# Biomass competition unifies individual and community scaling patterns

**DOI:** 10.1101/2024.03.25.586593

**Authors:** Lorenzo Fant, Giulia Ghedini

## Abstract

Both metabolism and growth scale sublinearly with body mass for most species. Ecosystems show the same sublinear scaling between production and total biomass but ecological theory cannot reconcile the existence of these nearly identical scalings at different levels of biological organization. We solve this paradox using marine phytoplankton to connect individual and ecosystem scalings across three orders of magnitude in body size and biomass. Competitive interactions determined by biomass, rather than differences in species size, slow metabolism in a consistent fashion across species that dominates over species-specific peculiarities, resulting in a unique behavior across community compositions. The allometry of ecosystem production thus emerges from this metabolic density-dependence, independently of the equilibrium state or resource regime of the system. Our findings demonstrate the mechanistic basis of ecosystem allometries, unifying aspects of physiology and ecology to explain why growth patterns are so strikingly similar across scales.

## Introduction

Ecosystems show remarkable regularities which suggest that their functioning is bound by common organizing principles ^1–4^. One of the most striking patterns is the power law between biomass production and total biomass which follows a sublinear scaling (often a near ¾ power law) and is independent of the ecosystem considered ^3^. The recurrence of the same scaling across ecosystems implies that communities of different species function similarly, independently of species composition and interactions ^3,5^. However current theory cannot explain how these regular ecosystem-level patterns emerge and how they can be independent of species size ^6,7^.

Across most taxa, individual metabolism and growth scale allometrically with body mass following a power law with an exponent *α* < 1 ^8–10^. This sublinear scaling means that, while larger organisms have greater metabolic (production) rates in an absolute sense, they consume less energy per unit mass compared to smaller organisms (valid for any scaling < 1). Since ecosystem metabolism and production increase with total biomass at a similar rate (∼ ¾), metabolic theory would predict that this pattern results from changes in species size: ecosystems of larger biomass should be dominated by larger organisms that have lower metabolism per unit mass ^3^. However empirical data do not support this explanation as size structure is nearly invariant across ecosystems ^3,5^. An alternative hypothesis is that sublinear scaling in ecosystems is a consequence of density-dependent processes which slow production as biomass accumulates ^3,6,11^. But how these processes operate to control biomass growth in such a consistent way across species and ecosystems – positing a common underlying mechanism – remains unknown.

To solve this puzzle, we overcome a main limitation of current theory. It is often assumed that metabolism-size relationships of species in isolation hold for these same species in communities, which has little empirical support ^11–14^. Competition for resources alters energy use and many organisms reduce their respiration rate in denser populations ^15–21^. Here we provide formal account for how density-dependent processes affect individual metabolism, biomass production, and their scaling with body size. We explore both metabolism and production because, while they are correlated, it is unclear which one drives the other. By doing so we demonstrate that the effect of competition on organismal physiology is the key to reconcile individual and ecosystem scalings.

We base our assessment on marine eukaryotic phytoplankton, a system of global importance for primary production ^22,23^. The size diversity of this system allows us to explore scaling relationships across three orders of magnitude in biomass at different scales (individuals, populations, species pairs, communities) ^24^. To demonstrate the generality of our findings we use geographically separate and independent phytoplankton communities (Australia, AU ^18^ and Portugal, PT), measured under a total of five environmental conditions in the laboratory. In each location, we sourced five species from local culture collections and grew them alone (monocultures) or together (communities). In the first location (AU) we also tested all species pairs, and cultures were grown under ideal conditions of light and salinity. In the second location (PT) cultures were grown under four combinations of decreasing light and salinity, creating a range of suboptimal environments. All cultures were initiated from a small biovolume of phytoplankton (a proxy for biomass, obtained as the product of cell volume and cell number); we then tracked changes in cell size, biovolume density, growth, and metabolism (photosynthesis, respiration) from exponential growth to stationary phase.

## Results

### Production and metabolism scale sublinearly from organisms to communities

Our phytoplankton communities display the same sublinear scaling observed across terrestrial and aquatic ecosystems ^1,3^. We find evidence of this power law scaling for production (i.e. biovolume growth, *α* = 0.79, confidence intervals CI: 0.77, 0.81, Fig. 1B), respiration (*α* = 0.74, CI: 0.68, 0.80, Fig. 1C), and photosynthesis (*α* = 0.61, CI: 0.53, 0.69, Supplementary Information Fig. S1) with total biovolume. Here and for the remaining of the manuscript, we separate data according to geographic origin (AU, PT) and light conditions (for the PT data), ignoring salinity manipulations because they have no appreciable effect on the trends we observe, although they might partly explain the increased variability we found in the second dataset.

**Fig. 1.**
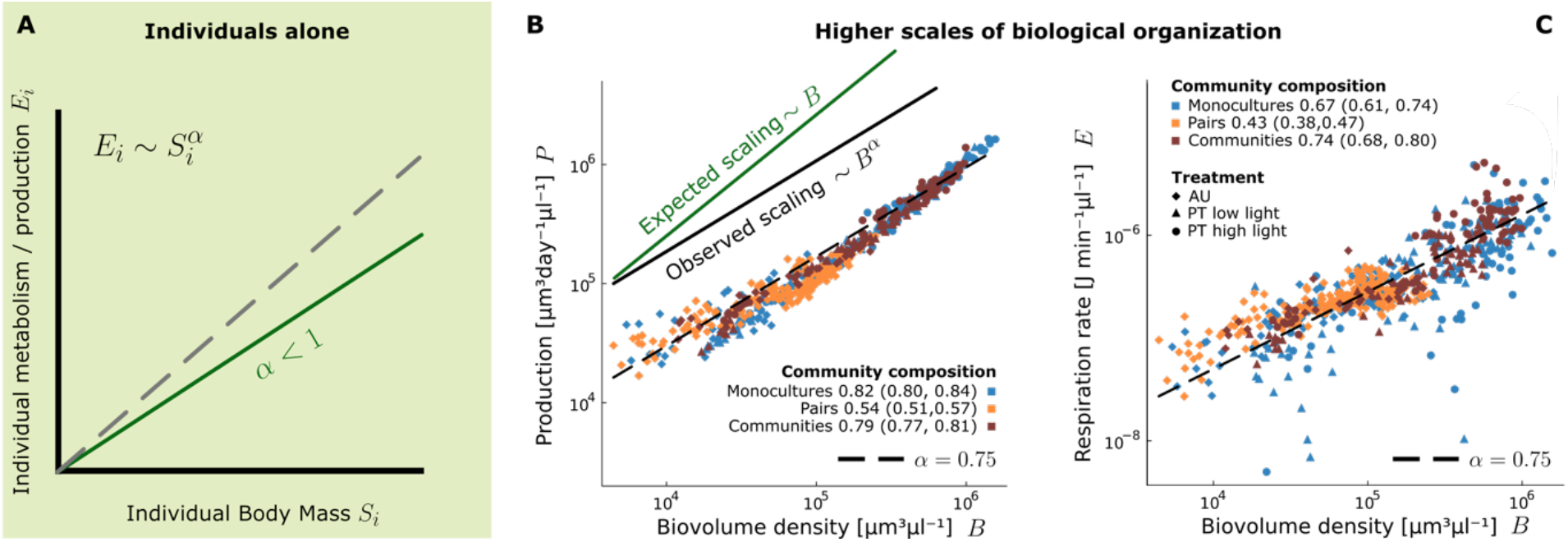
Production and metabolism scale allometrically with a nearly identical exponent across scales, in contrast with predictions from individual scaling data. (**A**) Individual metabolism and production scale hypo-allometrically (*α* < 1) with body mass across most taxa: 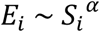. (**B**) According to this sublinear scaling at the individual level, the total production (or respiration rate) of a population of organisms of similar size should scale linearly with total population biomass (biovolume): 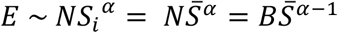 (expected scaling *∼ B*) because the total number of organisms 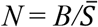. Contrary to this prediction, production (B) and respiration rate (**C**) scale sublinearly with biovolume across all monocultures, species pairs, and communities even if species size varies by three orders of magnitude.

We provide direct empirical evidence of the mismatch between theory and empirical patterns by measuring scalings at different biological levels of organization for the same system. The scaling of production or metabolism with biomass is remarkably consistent as we move from populations of different species (monocultures), pairs of species, and communities (Fig. 1, Fig. S1). The same sublinear scaling holds independently of species composition and with minor effects of environmental conditions, which we explore later. The regularity of these scalings is surprising because we see no effect of individual size on population or community metabolism. Individual metabolism scales non-linearly with size (Fig. S2) ^23^, so two systems (communities or populations) of equal biomass density but different size composition should not have the same metabolism: a system composed by smaller organisms should respire more than a system of larger organisms if the individual scaling is hypo-allometric (*α* < 1; and the other way around if *α* > 1). Let us consider individual populations (monocultures) where all organisms have similar sizes. Total population metabolism should scale linearly with total biovolume and with a size-dependent slope (intercept on a log-log scale). Contrary to this prediction, metabolism increases with biomass with the same sublinear scaling for all monocultures (*α* = 0.67, CI: 0.61, 0.74, Fig. 1C), even when our species vary in size by three orders of magnitude. The same holds for production (Fig. 1B).

The regularity of these relationships implies that, no matter the scale of organization and species composition, the total metabolism (or biomass production) of a system can be predicted from its total biovolume alone, both at and far from equilibrium. This is surprising because the only condition under which total biomass predicts total metabolism (independently of composition) is when all organisms have the same metabolic rate per unit mass – that is when metabolism scales isometrically with body size ^11^.

### Resolving the paradox: the coexistence of allometric and isometric scaling

Here we demonstrate, first mathematically and then empirically, that allometric and isometric scalings can coexist because they pertain to different comparisons: traditional comparisons across species, which focus on intrinsic species traits in the absence of density dependence (that yield allometry), and comparisons of interacting individuals embedded in an environment with the same biomass density (that yield an isometric scaling). We base our reasoning on respiration but we demonstrate below that the same holds for production.

Considering systems at a fixed biomass density *B*^*^, the number of organisms *N* is inversely proportional to their average size 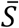 so that 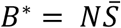 ^25^. The individual metabolism *E*_*i*_ is the total metabolism *E* divided by the number of individuals *N*. As qualitatively observed in Fig. 1, total metabolism scales with total biomass as *E* ∼ *B*^α^ across communities, independently of the composition of the system and the average size 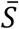 of its components ^1,3,11^. These considerations inevitably lead to a linear relation between size and individual metabolic rate when considering samples at constant biovolume:

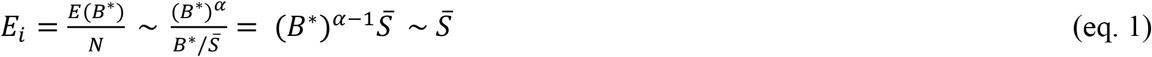

This result is independent of the scaling observed between total metabolism and total biomass (the value of *α*). Community metabolic scaling could follow any trend but, as long as it is not affected by the composition of the community (i.e. metabolism scales with biomass in the same way for all communities), it still leads to the isometric scaling of individual metabolism with size (Fig. 2A), and *vice versa*.

**Fig. 2.**
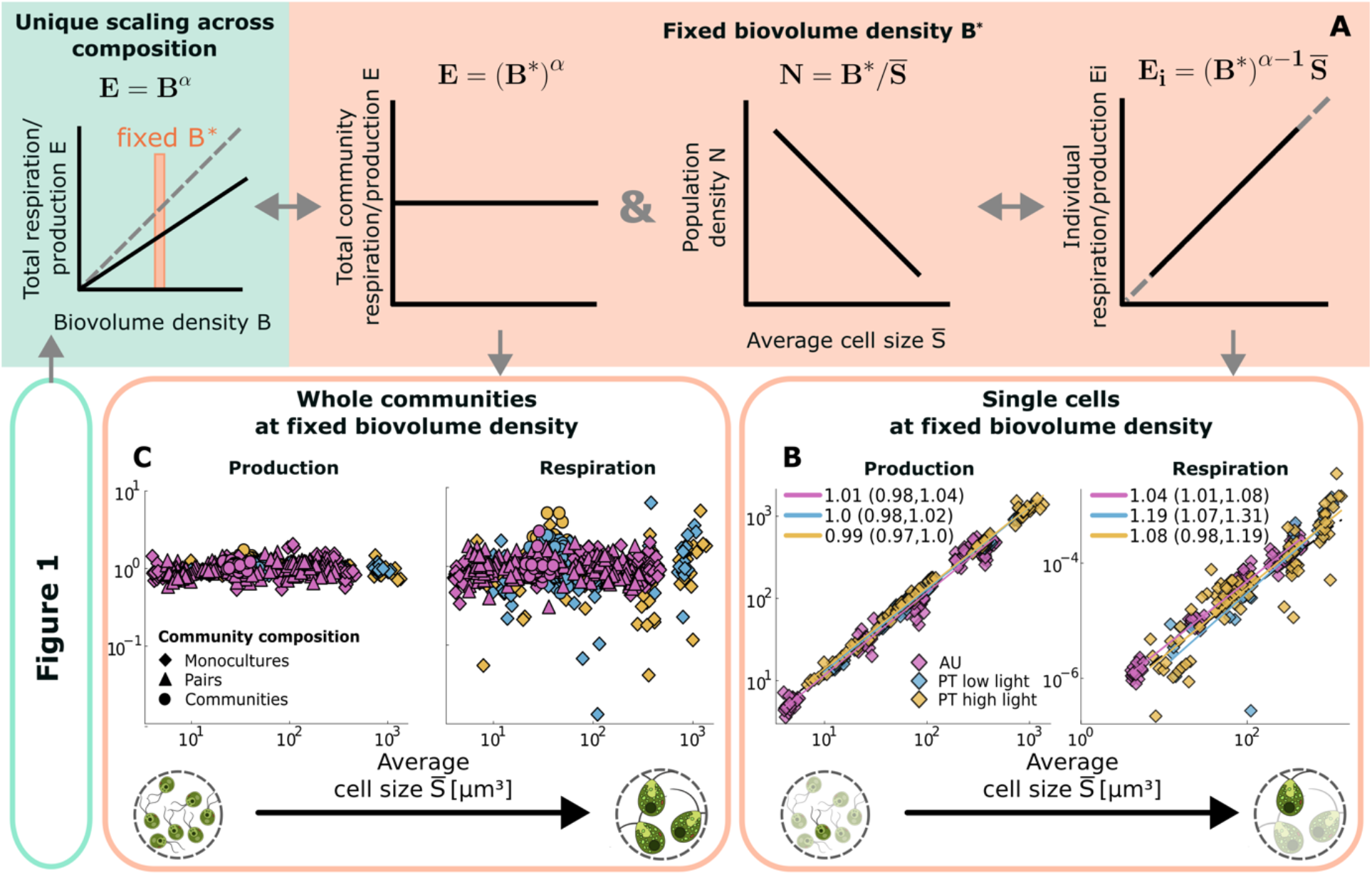
Theoretical predictions and empirical validation of the isometric scaling between individual metabolism and size in systems at fixed biomass density. (**A**) If there is a unique scaling between total metabolism and total biomass density as observed in Fig. 1 (i.e. the scaling is not affected by community composition; green), individual metabolism should scale isometrically with size when comparing systems at fixed biomass density. The same should hold for production. (**B**) We confirm this result empirically: *per capita* respiration (J/min) and production rates (μm^3^/day) scale linearly with cell size across phytoplankton species and environments when comparing organisms at fixed biovolume density (see note about low light environment in Supplementary Information). The broken black line has a slope of 1 for comparison. Here we only use data from monocultures but the scaling is robust even when considering data from communities (Fig. S5). (**C**) As a consequence of isometric scaling, organismal size has no effect on total energy use or production rates when we compare systems at equal biomass density. Whole population/community plots show normalized data, divided by the average respiration (production), as explained in Methods.

Isometric scaling is only valid at fixed biomass densities. Considering instead systems at fixed population densities (number of organisms *N* = *N*^*^, which includes traditional estimates of *per capita* metabolic scaling based on *N*^*^ = 1), the average size becomes just a proxy for total biomass; thus, when evaluating the individual metabolic rate as a function of size, one retrieves the initial scaling of metabolism to biovolume (Fig. S3):

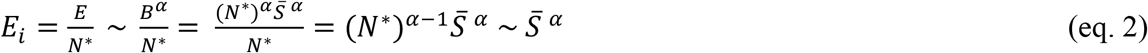

If neither biomass nor the number of organisms are fixed quantities, metabolism can scale with size in several ways (Fig. S3). Since metabolism-size relationships are often estimated for different organisms in varying conditions of population or biomass density, this could partly explain the variability in the scaling exponents reported in the literature ^8,9,23,26^.

Therefore, if the qualitative observation of Fig. 1 is correct (i.e. total metabolism scales with biomass independently of composition), competition with an equal amount of biomass must shift the scaling of *per capita* metabolism with body size from allometric (eq. 2) to isometric (eq. 1).

We validate these theoretical predictions with empirical data by testing the linearity between individual metabolic and production rates with cell size at fixed biomass density. We use the full range of biovolume of our system by rescaling all data points to a single biovolume value in the centre of the range. As predicted, rescaling individual metabolism or production to a fixed biovolume unravels an isometric scaling between *per capita* rates and cell size (Fig. 2B respiration and production, Fig. S4A photosynthesis). This result also holds for the *per capita* metabolic rates of “average*”* individuals in mixtures of species (Fig. S5). By demonstrating this theoretical result, we quantitatively validate our observation: community metabolism and production are independent of species composition and average size (Fig. 2C respiration and production, Fig. S4B photosynthesis). We further show that individual metabolism scales non-linearly with cell size when comparing systems at equivalent population densities according to eq. 2 (Fig. S6).

The observed shift in scaling from allometric (when comparing individuals of different sizes at fixed population densities) to isometric (when individuals of different sizes compete with an equal amount of biomass) implies that the sublinear scaling of metabolism with mass is driven by the increase in biovolume – in other words, metabolism per unit mass declines with increasing biovolume and this decline is the same regardless of how biovolume is distributed (i.e. if it surrounds the organism or if it is part of it, as shown in Fig. 2B). Indeed, when we remove differences in biovolume and compare small and large phytoplankton cells competing with the same amount of total biovolume, their metabolism scales isometrically with size.

### Everybody is anybody: composition does not affect metabolic density-dependence

Now that we observed that biomass competition affects metabolism and production in the same way across individual species, we investigate at a finer scale whether there are smaller species-specific effects when these species interact directly in communities. Specifically, we use our data from phytoplankton communities to test the importance of the following two factors (Fig. 3A):

**Fig. 3.**
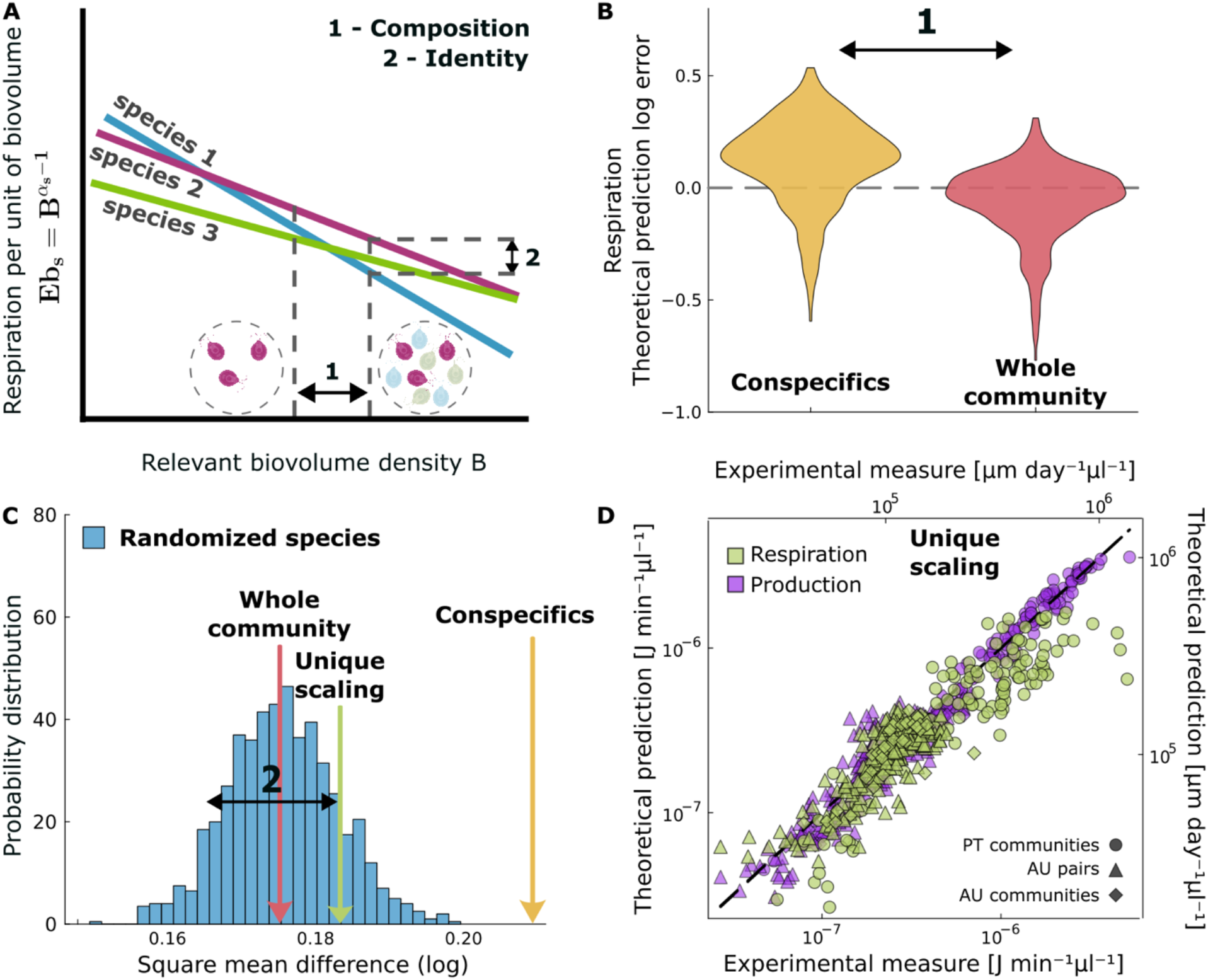
Predictions of community metabolism based on different models of metabolic density-dependence. (**A**) Schematic showing that metabolism per unit biovolume might decline with increasing biovolume density at different rates for each species. We use these relationships, based on monoculture data, to test the importance of two factors in driving community metabolism: 1) *competitors’ composition* (are metabolic declines driven equally by intra-inter-specific competitors?), 2) *metabolic identity* (do species-specific differences considerably affect community rates?). (**B**) Error of the predictions testing factor 1. We can correctly estimate community metabolic rates if we consider the total biovolume of the community. If we account conspecifics only, we overestimate community rates. (**C**) Species identity (factor 2) does not significantly affect community respiration. If we randomize the association between species-specific declines in metabolism and the biovolume of species in the community, we obtain a distribution of estimates (blue) that contains the prediction made using the correct association (whole community, magenta). Predictions based on conspecifics are outside of this range. (**D**) Since species identity has minimal effects, we can estimate total community metabolism (green) or production (purple) using a unique scaling between metabolism (production) per unit mass and total biomass across all species (green arrow in panel C, refers to metabolism). See Fig. S10 for production in Supplementary Information.

1. *competitors’ composition*: the differential impact of species on one another. Specifically, we investigate whether conspecifics exert a more pronounced influence on a species metabolism compared to individuals of other species (interspecifics);
2. *metabolic identity*: the importance of metabolic differences among species, seeking to understand if distinct species exhibit varied responses to competition.

We start by partitioning the relative importance of intra- and inter-specific competitors on species’ metabolism (1), assuming that species identity matters (2). Metabolism is density-dependent but the rates of these declines might be species-specific (Fig. 3A). We calculated these rates for each of our species using monoculture data and used these species-specific relationships to calculate the metabolism per unit biovolume of each species in the community, according to two extreme hypotheses:

a. competition for resources is stronger within species than among species ^27^, so only conspecifics reduce metabolism while interspecific competitors have negligible effects; hence metabolism per unit biovolume declines only in response to the *biomass density of conspecifics*: 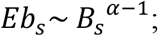
b. phytoplankton compete for similar resources, so intra- and inter-specific competitors have equal effects on metabolism; hence metabolism per unit biovolume declines in response to the *total biovolume density* of the community: 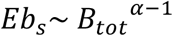.

Using either approach, we estimated the metabolism of each species in the community knowing its biovolume, and finally the total community metabolic rate as the sum between species (see Methods for details). We compare these predictions with the metabolic rates measured experimentally on communities throughout their growth. Predictions based on the total community biovolume are accurate (Fig. 3B). Conversely, if we do not account for interspecific competitors, we overestimate community metabolic rates because we underestimate the level of metabolic suppression driven by competition. We also explored an intermediate situation in which interspecifics might affect metabolism in a weaker way than conspecifics by measuring competition effects among all pairs (this test was only possible for AU data which included all species pairs). Despite its greater specificity, this approach does not improve predictions (Fig. S7-S9). Thus, metabolism declines identically in response to intra- and inter-specific competitors.

Now we challenge our second assumption: does species identity matter? We find that it does not. If we randomize the association between species-specific declines in metabolism and the biovolume of species in the community, we obtain a distribution of estimates that contains the prediction made before, using the correct association (Fig. 3C); in other words, predictions based on randomised associations perform similarly to species-specific predictions. The variability in metabolic declines with biomass between species therefore does not meaningfully affect community outcomes, at least based on our experimental accuracy. Importantly, the randomized distribution does not contain the prediction based on conspecifics biovolume (hypothesis a), confirming that the biovolume of all competitors is the quantity that affects species metabolism in communities.

If species identity has weak effects, then we can ignore it and estimate a unique relationship between metabolism and biomass by merging all measures obtained across monocultures (in each environment):

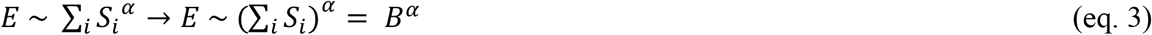

Despite the simplicity of this approach based on a unique decline in metabolism with biomass across species, we can correctly predict community metabolic rates in all environments and across all growth phases (Fig. 3D). This approach performs worse than that based on species-specific rates (*α*whole community*α*) but is still within the randomized distribution. So we cannot state that there is no variability in metabolic responses between species, but this variation is not sufficiently strong to prevent us from estimating community rates using a unique metabolic decline across species (Fig. 3C and D; SI for details on the importance of species identity). Thus, even in a system of interacting species, metabolism slows with increasing biomass at the same rate on average among species, regardless of the nature of the biomass (i.e. relative abundance of intra- and inter-specifics, and their size).

All the considerations we have done for respiration extend to biomass production which is also invariant with respect to species size and composition (Fig. S10). These conclusions, instead, only partially hold for photosynthesis. As for respiration, we can correctly predict community photosynthesis by estimating the decline in species rates in response to the total biomass of the community (rather than conspecific biomass) (Fig. S11B). But we observe a stronger effect of species identity (Fig. S11C), which might be partly due to the unimodal scaling of photosynthesis with cell size (Fig. S4B, phytoplankton species of intermediate size have higher rates of photosynthesis per unit mass) ^28^ and to the fact that species-specific responses to environmental conditions (light availability) might be more pronounced for photosynthesis than respiration.

### Metabolism mirrors production scaling across environments

So far we observed that metabolism and production scale similarly with species size and total biomass. Nonetheless, our results do not require the two scalings to be the same as they do not rely on the specific values of *α*. In this last section we tie together the two by showing that there is a quantitative connection between metabolic suppression and the sublinear scaling of ecosystem production with total biomass. We make this link by focusing on the environment. As would be expected based on differences in resource availability, the environment (primarily differences in light availability) influences the scaling between total metabolism and total biomass but this variation is not random: the biomass-dependence of respiration rates quantitatively tracks declines in biomass growth within each condition and this tight relationship persists across scales of organization (from populations to communities) (Fig. 4), despite marked differences in environmental conditions and resource availability in our cultures (Fig. S12). Notably, the quantitative link between metabolism and production is not visible for all energy fluxes, suggesting that respiration captures metabolic processes associated with growth more closely than photosynthesis (Fig. S13). While we cannot establish that metabolism governs production and not the other way around ^29^, our data show that biomass competition slows metabolism in a very similar and predictable fashion across species, constraining growth not only in individual populations ^16^ but also in entire communities and throughout their whole growth process (also when far from equilibrium).

**Fig. 4.**
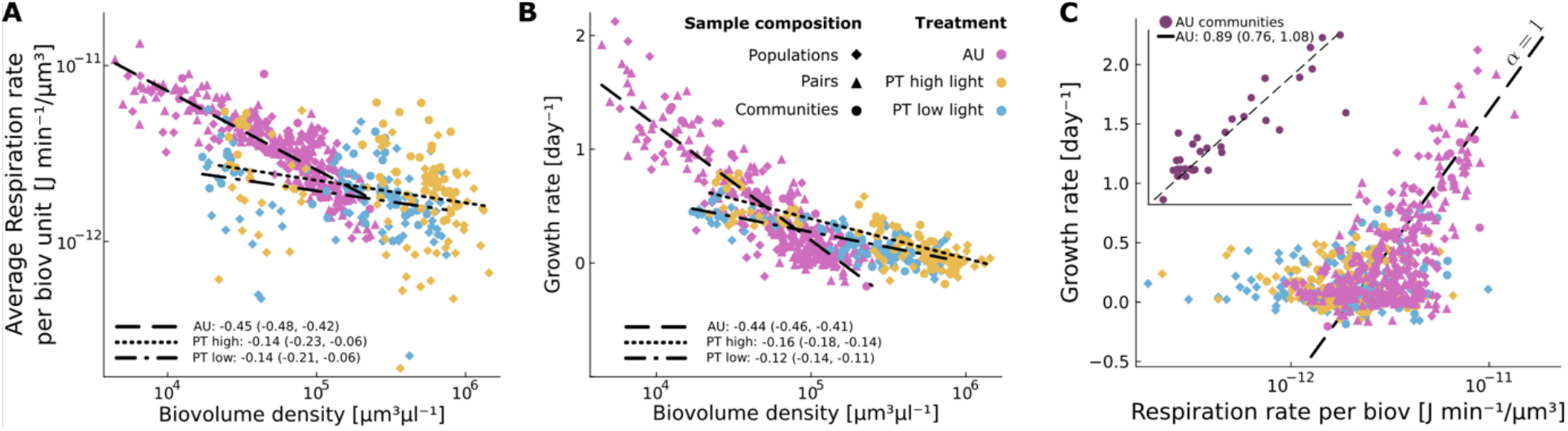
Declines in metabolism with biomass track declines in production across environments. (**A**) Respiration rates per unit biovolume (calculated as the geometric mean between consecutive measurements) decline with total biovolume density at different rates depending on the environment. (**B**) In each environment, metabolic declines quantitatively track reductions in biomass growth, not only in populations but also in communities – indicating a strong level of community regulation on both metabolism and growth (**C**).

## Discussion

Ecosystem productivity scales predictably with total biomass, independently of species size and composition ^3^. These size-independent patterns seem incompatible with the allometry of growth and metabolism observed at the individual level for most taxa ^5^. We demonstrate the connection between individual and ecosystem scalings by showing that biomass competition influences organismal metabolism identically across species that compete for similar resources. While species of different sizes grow and respire at different rates per unit mass when compared at equal population densities (focusing on body size properties, i.e. classic individual allometries), the allometry between individual metabolic rate and body size collapses onto isometric scaling once we account for changes in metabolism in response to biomass competition, both within and between species. Thus, competition with an equal amount of biomass alters the scaling of metabolism with size in a defined and predictable way that holds across growth phases and environments, and is independent of the composition of the biomass. This result solves some of the inconsistencies and variability in metabolic scalings ^19,30–34^ [reviewed in ^8,23^], and shows how essential it is to account for varying levels of competition when estimating scaling exponents.

We do not know the specific mechanism that leads to consistent changes in metabolism across species. Ecosystems display allometric patterns of resource transport efficiency that resemble size-dependent patterns in organismal metabolism ^1,4^. So the generalized metabolic decline we observe with biomass might emerge both because increases in biomass density alter the flow of resources according to common organizing principles ^1,4^ and because many species respond to changes in densities through behavioural and physiological adjustments that lead to metabolic suppression ^15,17,19–21,35,36^. The resulting scaling patterns might thus be independent of the specific nature of interactions, at least when species compete for similar resources ^6,37^. We offer the first empirical demonstration of this hypothesis ^6^ and show that community functioning is tightly integrated – to the point that extending the relationship between metabolism and mass from organisms to entire communities gives a reliable representation of community functioning.

## Materials and Methods

### Experimental set ups

We combined two geographically distinct datasets of marine phytoplankton. Both datasets used species of marine phytoplankton obtained from culture collections; species were cultured in temperature-controlled rooms at 22 ± 1°C using autoclaved Guillard’s f/2 medium, prepared with filtered natural seawater.

The first dataset (AU) is from Ghedini et al. (2022) ^18^ where they grew five species of marine phytoplankton in three species diversity treatments over 10 days: monoculture, in pairs or communities with all five species. Each monoculture and species pair were replicated three times and communities five times (N = 50 cultures). The work was performed at Monash University, Australia, and the species used were obtained from the Australian National Algae Culture Collection: *Amphidinium carterae* (CS-740), *Tetraselmis* sp. (CS-91), *Dunaliella tertiolecta* (CS-14), *Tisochrysis lutea* (CS-177) and *Synechococcus* sp. (CS-94). All cultures were placed in cell culture flasks filled to 100 ml and grown on a 14–10 hr light– dark cycle under non-saturating irradiance levels (115 μmol m^−2^ s^−1^) at ambient salinity (35 ppt). Flasks were shaken and randomly rearranged on the shelves every day. Nutrients were added daily by replacing 10% of medium from each flask with fresh f/2 medium. All cultures were started with an initial total biovolume ∼ 6 × 10^8^ μm^3^, where biovolume is the product of cell size (volume) and number of cells and is used a proxy for biomass in phytoplankton. Cultures were sampled as detailed below on each day for the first five days and at alternate days afterwards for a total of eight sampling times (day 0, 1, 2, 3, 4, 6, 8 and 10).

The second dataset (PT, unpublished) was collected at the Gulbenkian Science Institute (IGC) in Portugal using five phytoplankton species obtained from the Roscoff Culture Collection: *Amphidinium carterae* (RCC88), *Dunaliella tertiolecta* (RCC6), *Phaeodactylum tricornutum* (RCC2967), *Tisochrysis lutea* (RCC90), *Nannochloropsis granulata* (RCC438). These species were grown either in monoculture or in a community over 16 days under two levels of salinity (35 or 20 ppt) and light (60 or 30 μmol m^−2^ s^−1^) in cross combination to simulate a gradient of stressful environments. We set up 5 replicate communities (a mix of the five species in equal biovolumes) and 2 replicate monocultures of each species for each level of salinity and light in glass bottles filled to 200 ml (N = 60 cultures). The position of the cultures was randomized at each sampling day and cultures were bubbled continuously for mixing. We started with an initial total biovolume of ∼ 4 × 10^9^ μm^3^ for each treatment.

We tracked changes in the abundance, size, and biovolume of species over time through microscopy; concomitantly, we measured the metabolism of monocultures and communities using respirometry (photosynthesis and respiration). We maintained salinity treatments by adding small amounts of distilled water when needed. Communities and monocultures were sampled 7 and 6 times, respectively over the course of 16 days to measure changes in biovolume and metabolism as detailed below.

### Cell size, population and biovolume density

In both experiments, 1 ml of sample from each culture was fixed with 1% Lugol’s solution to quantify cell size and abundance. From these fixed samples, we loaded 10 μl onto a cell counting chamber (Neubauer improved) and we took photos of the sample with an Olympus IX73 inverted microscope using a 400x magnification. Photos were processed in Fiji/ImageJ ^38^ to quantify the cell volume (μm^3^), number of cells of each species (μl^−1^), and biovolume as their product (μm^3^ μl^−1^). Cell volume was calculated from the major and minor axis of each cell by assigning to each species an approximate geometric shape (prolate spheroid for all species, except *Synechococcus, Tisochrysis*, and *Nannochloropsis* for which we assumed a spherical shape) ^39^. The total biovolume of species mixtures (pairs or communities) was calculated as the sum of each individual species’ biovolume.

### Metabolism

Photosynthesis and respiration rates were measured from changes in percentage oxygen saturation under light (photosynthesis) or dark conditions (respiration) using 24-channel sensor dish readers (SDR; PreSens Precision Sensing GmbH). Measurements were done on 5 ml (AU) or 2 ml (PT) glass vials with integrated oxygen sensors.

The system was calibrated with 100% and 0% oxygenated water prior to each experiment. We quantified photosynthesis as the rate of oxygen production under the same light intensity at which the cultures were grown, over a period of 30 minutes or less if cultures approached 250% earlier (the maximum value the instrument can read). Respiration was quantified as the rate of oxygen decline over 30 minutes following light exposure. We added a 2% solution of sodium bicarbonate to each vial to avoid carbon limitation during photosynthesis. We added blanks prepared with the supernatant of the samples on each SDR reader to account for drift and bacterial respiration (12 and 24 for the two datasets respectively).

In both experiments, the rate of photosynthesis or respiration of the whole sample (VO_2_; units μmol O_2_/min) was measured as VO_2_ = 1 × (( m_a_m_b_)/100 × VβO_2_) following ^40^, where m_a_ is the rate of change of O_2_ saturation in each sample (min^−1^), m_b_ is the mean O_2_ saturation across all blanks (min^−1^), V is the sample volume in litres and VβO_2_ is the oxygen capacity of air-saturated seawater for the temperature and salinity of the sample (μmol O_2_/L). The first three minutes of measurements were discarded for all samples for photosynthesis and respiration rates were calculated after 10 minutes of dark when rates showed a linear decline. Rates of photosynthesis and respiration (μmol O_2_/min) were converted to calorific energy (J/min) using the conversion factor of 0.512 J/μmol O_2_ ^41^ to estimate energy production and energy consumption respectively as in previous work ^42^.

### Data analysis

Data were analysed and visualised through a *Julia* pipeline.

1. *Data filtering:* We discard negative respiration measures. Negative values of respiration are obtained when the slope of the blanks in steeper than that of the sample, which indicates either some error in the preparation/seal of the vial or that the sample does not contain enough (live) phytoplankton biomass to differentiate their respiration from that of bacteria (blanks). The number of discarded samples is: 12/197 (PT monocultures), 11/279 (PT communities), and none in the other dataset: 0/105 (AU monocultures), 0/210 (AU pairs), 0/35 (AU communities).
2. *Data normalisation*: Respiration rates, photosynthesis rates, total biovolume values are normalised by the sample volume (2 ml for PT data, 5 ml for AU data).
3. *Community/population metabolic rate to total biovolume*: We fit linearly the logarithm of respiration (photosynthesis) rates as a function of the logarithm of biovolume density. Thus we do not assume that there is an intrinsic power law relationship between these two quantities. In fact, the growth in biovolume we observe for each replicate is smaller than two orders of magnitude, not allowing to establish a reliable functional form. Nonetheless, the log-log relationship allows us to find a functional form enabling the following analysis. The fits were done by grouping data in two ways: 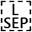 a – by environment (AU, PT high light, PT low light) – we find no significant effects of salinity manipulations in the PT data, thus we do not consider this factor; b – by environment and species. By comparing the predictions of community respiration rates obtained with the fits of the two groupings, we will show that nor the species identity nor the competitors composition plays a relevant role in determining total respiration rates.
4. *Respiration rate per unit of biovolume:* We divide the measured respiration rates by the total biovolume. We then fit linearly the logarithm of rates per unit of biovolume as a function of the biovolume logarithm. Here we fit the data by grouping, as above, for (a) environment only (b) species and environment.
5. *Individual metabolic rate to population biovolume*: We calculate the average individual respiration (photosynthesis) rates (per cell) by dividing the total respiration rates by the cell density. We then fit linearly the logarithm of individual rates as a function of the biovolume logarithm. Here we fit the data by grouping according to species and environment as individual metabolic rates depend on the size of individuals themselves.
6. *Community/population metabolic rate to cell density*: Similarly to above, we fit linearly the logarithm of respiration (photosynthesis) rates as a function of the logarithm of cell density (number of cells).
7. *Fitting method*: We use ordinary least squares method (OLS) for fits on log-transformed data, which is recommended when the error on the x-axis (biovolume) is smaller than the error on the y-axis (metabolism) and consistent with previous work (*3*).
8. *Rescaled metabolic rate at fixed biovolume or cell density*: Here we want to evaluate the slope of metabolism-size relationships when species are at the same biovolume or cell density. To do this, we use the monoculture data to rescale the respiration (photosynthesis) rate of each species (at the population-level) to the value predicted at a fixed biovolume, using the scaling obtained in 3.b (environment + species). In this way every species was rescaled independently as 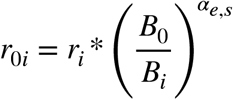, where *r*_*i*_ is the measured respiration rate, *B*_*i*_ is the measured total biovolume, *B*_0_ is the target (fixed) biovolume density (10^5^ μm^3^μl^-1^), and *α*_*e,s*_ is the exponent of the power law fitted at fixed environment *e* and species *s*. Such rescaling keeps the spread of the data on the y-axis (respiration rates) intact while removing variation on the x-axis (biovolume). The same procedure can be performed to fix the cell density to a value *N*_0_ by 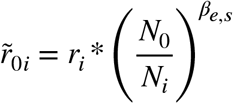 where *β*_*e,s*_ is the exponent found by fitting metabolism as a function of cell density.
9. *Average individual respiration rate:* by rescaling the total biovolume to a fixed value we lost the information on the total number of individuals. We thus use the relationship between the total biovolume *B*_0_ and the number of individuals *B*_0_ = *N*_0*i*_ * ⟨*B*_*i*_⟩ to calculate the number of individuals at the fixed biovolume. We know both *B*_0_ and the average size of each species ⟨*B*_*i*_ ⟩, thus the number of individuals is 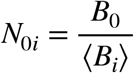. We then divide the community respiration rates at fixed biovolume by *N*_0*i*_ to calculate individual respiration rates.
10. *Predict community rates from monoculture data:* We use the relationship between *Respiration rate per unit of biovolume* and population biovolume density obtained for monocultures (point 3) to predict the metabolism of biovolume in each community with three methods:
  a. *Conspecifics*: we use the fits defined in 3.b (environment and species) to calculate the metabolic rate of biovolume for each species and in each environment from the biovolume density of conspecifics present in the community at each timepoint. This approach assumes that the metabolism of a species is only responsive to the presence of conspecifics, while other species have no effect. We then sum over all species to find community total metabolism;
  b. *Whole community*: we use the fits defined in 3.b (environment and species) to calculate the metabolic rate of biovolume for each species and in each environment from the total biovolume density of the community at each timepoint. This approach assumes that the metabolism of a species is equally affected by competitors, independently of their nature – it doesn’t matter who your competitors are, only how much biomass density surrounds you (see also *13*). We then sum over all species to find community total metabolism;
  c. *Unique scaling*: a unique (not species-specific) relationship between metabolism per unit of biovolume and biovolume that varies between environments; we calculate it using the grouping defined in 3.a (environment only). This approach assumes that the metabolism of a species is affected by biovolume in a way that is species independent. Identity does not matter; competition affects everyone in the same way.
  d. *Pairs:* we calculate the effect of intraspecific competition in species pairs by fitting the difference between the expected metabolism per unit of biovolume of each species (based on monoculture data) 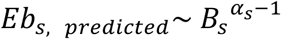 and the measured average metabolism per unit of biovolume (measured on pairs) 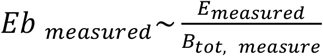. Specifically, we fit linearly the distribution of points with coordinates:

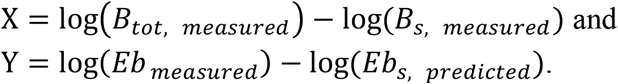

In this way we can estimate the effect of each species on each other obtaining the slopes β_*s,i*_ that express the effect of the species *i* on species *s*. The obtained values are used to estimate the effect of interspecific competition in communities as

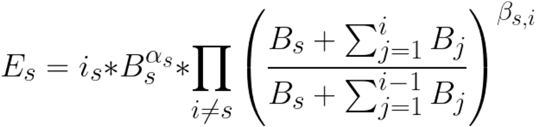

where *i*_*s*_ is the exponential of the intercept obtained in 3.b.
11. *Average respiration rate per unit of biovolume:* We calculate the geometric mean between consecutive measurements to have a quantity relatable to the average growth rate between consecutive measurements.
12. *Prediction error:* We calculated the difference (delta) between the estimates obtained above and the observations of community metabolism on a log-scale to visualise the offset of the prediction for each hypothesis.
13. *Growth rate:* Calculated as 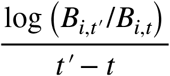. Where *B*_*i,t*_ and 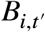 are two total biovolume measurements performed on sample *i* at time *t* and *t ‘* − *t*. For the AU experiment we calculated growth based on the 90% of the previous biomass because each sampling day we removed 10% of the sample.
14. *Productivity:* Calculated as 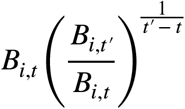 is the estimate for the daily biovolume increase of sample *i* at time *t*.

## Authors contributions

LF and GG conceived the idea of this manuscript. GG designed and performed the experiments. LF analysed the data with feedback from GG. GG drafted the manuscript and both authors contributed to the final version.

## Acknowledgments

We are grateful to Michel Loreau, Jacopo Grilli, Onofrio Mazzarisi, and Isabel Gordo for their comments on an earlier version of this manuscript. We thank Moritz Klassen for help with data collection. LF was supported by a postdoctoral PONTE fellowship from the Gulbenkian Science Institute. GG was supported by a fellowship from ‘‘la Caixa’’ Foundation (LCF/BQ/PI21/11830001; ID 100010434) and the European Union’s Horizon 2020 research and innovation programme under the Marie Sklodowska Curie grant agreement no. 847648.

