## Supplementary Information for "Biomass competition unifies individual and community scaling patterns"

**The PDF file includes:**

Supplementary Text
Figs. S1 to S13

**Supplementary Text**

*Supplementary text about Fig. 2B*

The low light environment deviates from this prediction showing hyper-allometric scaling; we investigated this deviation and found that it is driven by a species (*Phaeodactylum*) in the most extreme environment (combination of low light and low salinity where scaling is 1.28, CI: 1.05-1.51) probably indicating physiological stress. If we remove this extreme treatment, the scaling at low light also overlaps isometry (1.11, CI: 1.0-1.23).

*Species Identity importance*

Given the results obtained in Figs. 3 (main), S10 and S11, we stated that species identity is marginally relevant to determine the metabolism and production of communities. In fact, the predictions performed by correctly associating the scalings calculated in monocultures for each species to the biomass of the exact species in the community are as good as (respiration) or slightly better (production) than those obtained by randomly associating the biomasses to the scalings. The same is not true for photosynthesis, where species identity seems to matter. Moreover, given the intrinsic limited precision of experimental data, we cannot assess that there is no influence at all even for respiration.

Nonetheless, it is key to notice that such an influence would not affect the paper's main finding. Even if species identity were important, the general trend has been proved in Fig. 2 (main) and, even more importantly, the key concept at the foundation of our solution is that metabolic responses are determined by biomass competition, regardless of the competitors. The former was independently proved in Figs. 3B (main) and S7, S8, S9.

We think that the use of a unique scaling for all community $Eb \sim B_{tot}^{\alpha}$ is a useful tool to understand the general behaviour of our communities. If species identity will prove to be relevant it should be simply changed to $Eb \sim \sum_s i_s \, B_s (B_{tot})^{\alpha_s - 1}$ where $i_s$ is the exponential of the intercept obtained in Methods 3.b. Here too we can notice our key finding that respiration, production, and photosynthesis of biomass (and thus individuals) depend on the whole biomass density. Thus, in case exponents are sufficiently similar between species, as they proved to be, leads to the same findings of the paper.

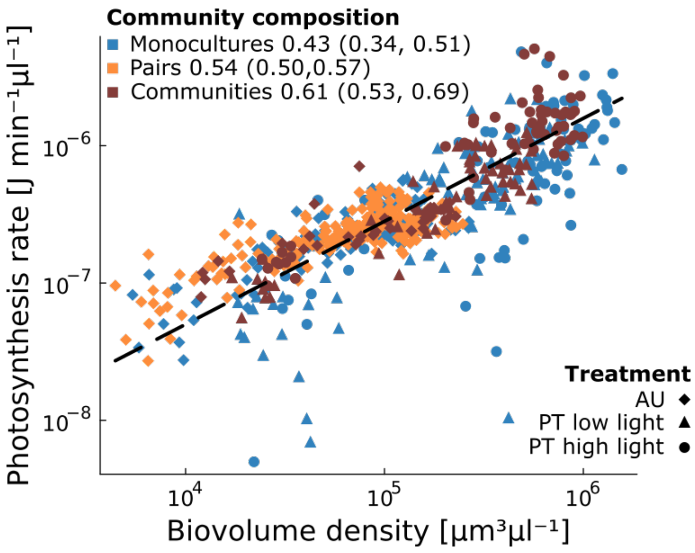


**Fig. S1. Photosynthesis scaling across levels of organization.**

Photosynthesis rate scales sublinearly ($\alpha < 1$) with biovolume density across different levels of organization, that span monocultures, species pairs, and communities in different environments and independently of species composition. The black line represents $\alpha = 0.75$ for comparison.

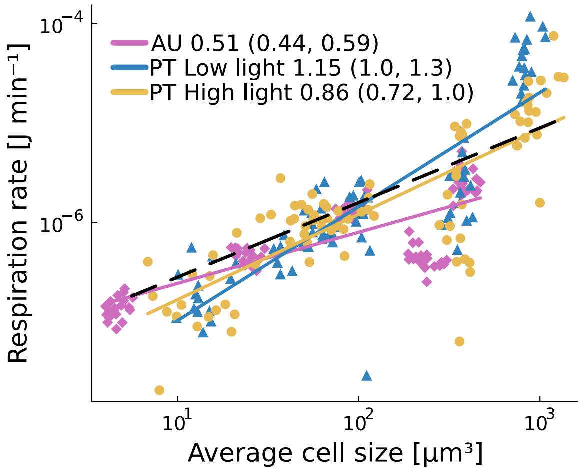


**Fig. S2. Scaling of *per capita* respiration rates with cell size.**

*Per capita* respiration rate scales non-linearly ($\alpha \neq 1$) with cell size for our phytoplankton species measured in monoculture. The black line represents $\alpha = 0.75$ for comparison.

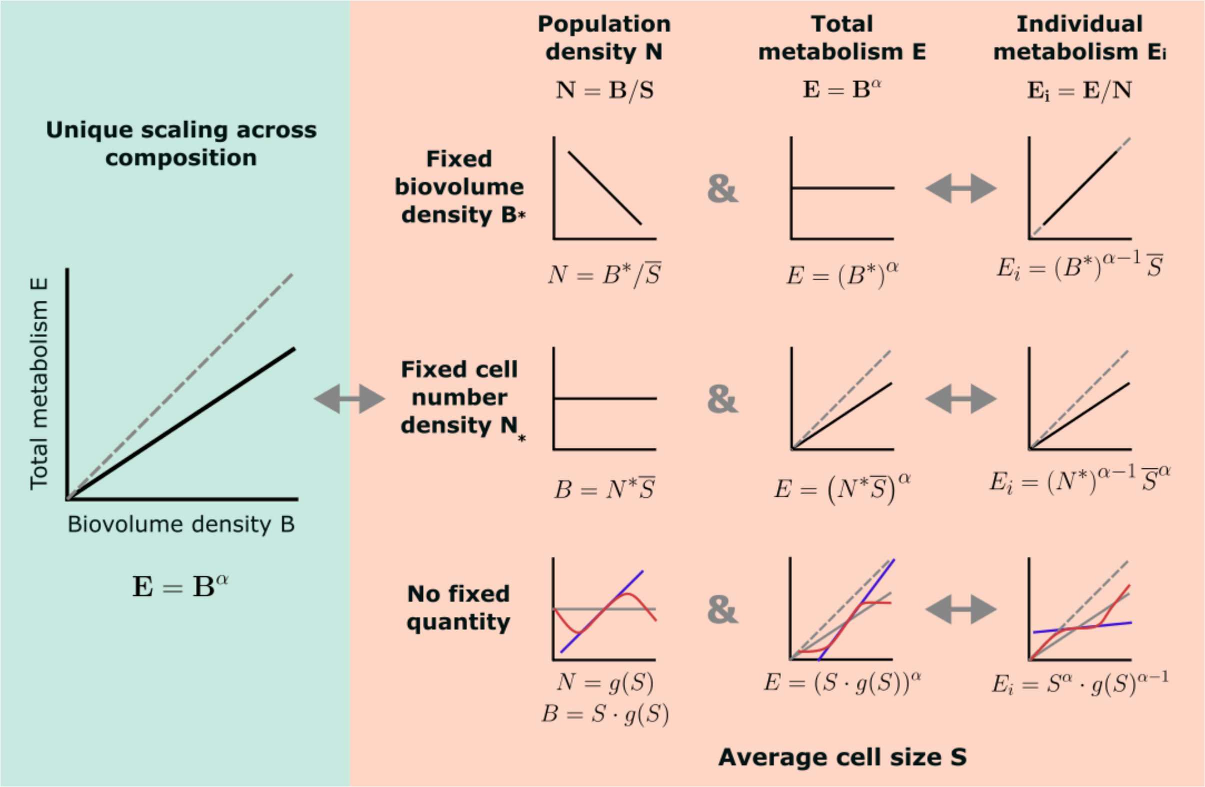


**Fig. S3**. **Schematic showing the implications of a unique scaling independent of composition for individual scaling patterns.**

If the scaling of total metabolism and total biomass is not affected by species composition (i.e. there is a unique scaling across communities, green), then when we compare systems at equal biomass density ($B = B^*$), *per capita* metabolism $Ei$ must scale isometrically with individual size (top row). If we consider instead systems at fixed population densities (number of organisms, $N = N^*$), the average size becomes a proxy for total biomass – thus, when evaluating the individual metabolic rate as a function of size, one retrieves the initial scaling of metabolism to biovolume ($\alpha$) (middle row). When neither biomass nor cell numbers are standardized, per capita metabolism can scale with size in several ways (bottom row).

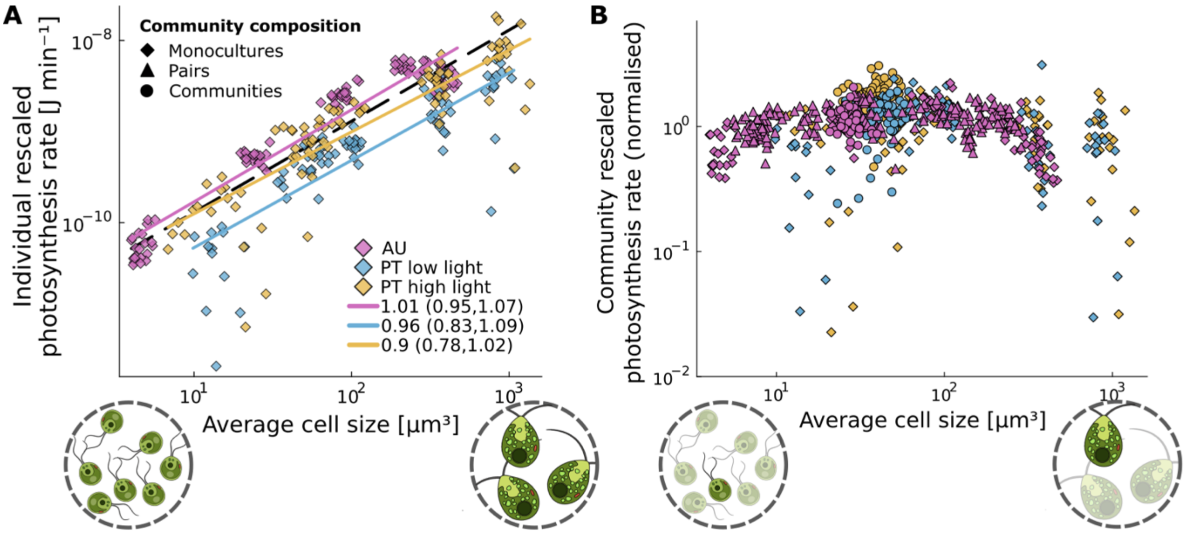


**Fig. S4. Scaling of *per capita* and total photosynthesis with cell size at fixed biovolume density.**

(**A**) *Per capita* photosynthesis rates scale isometrically with size across phytoplankton species and environments at fixed biovolume density. The broken line has a slope of 1 for comparison. The green circles represent individual phytoplankton cells; here we compare the photosynthesis rate of a small and large cell (dark green) surrounded by an equal amount of biovolume (pale green). (**B**) Organismal size has largely no effect on total energy use (here shown for photosynthesis) when we compare systems at equal biomass density, although total photosynthesis rates show a slight curvature (unimodal relationship) with size, in line with the fact that phytoplankton of intermediate size have higher biomass-specific metabolic rates.

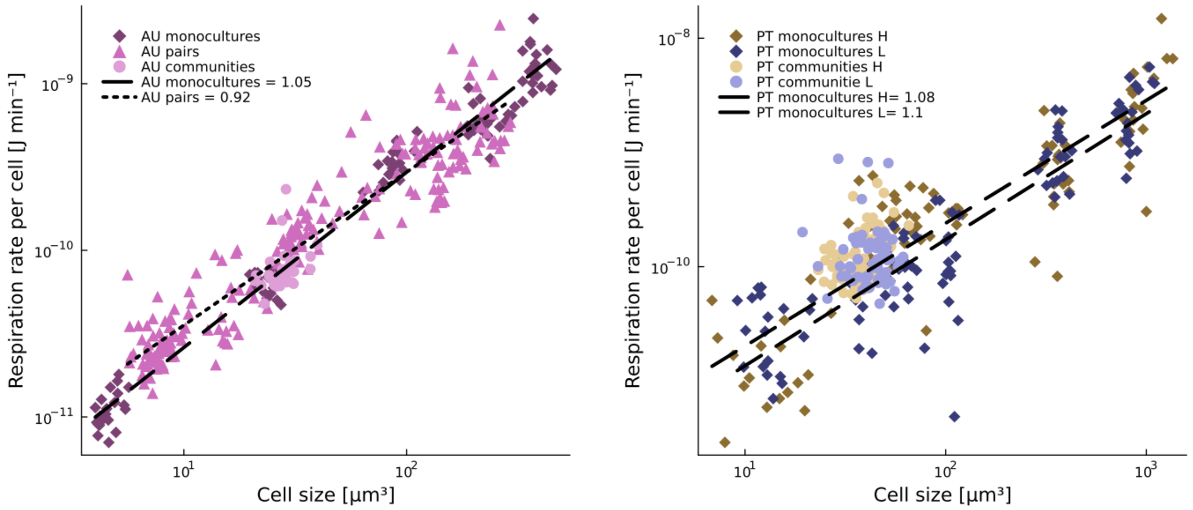


**Fig. S5. Scaling of *per capita* metabolism with cell size at fixed biovolume density including species pairs and communities.**

*Per capita* metabolism scales linearly with cell size at fixed biovolume, also when considering the average metabolism-average size scaling obtained from species pairs and communities.

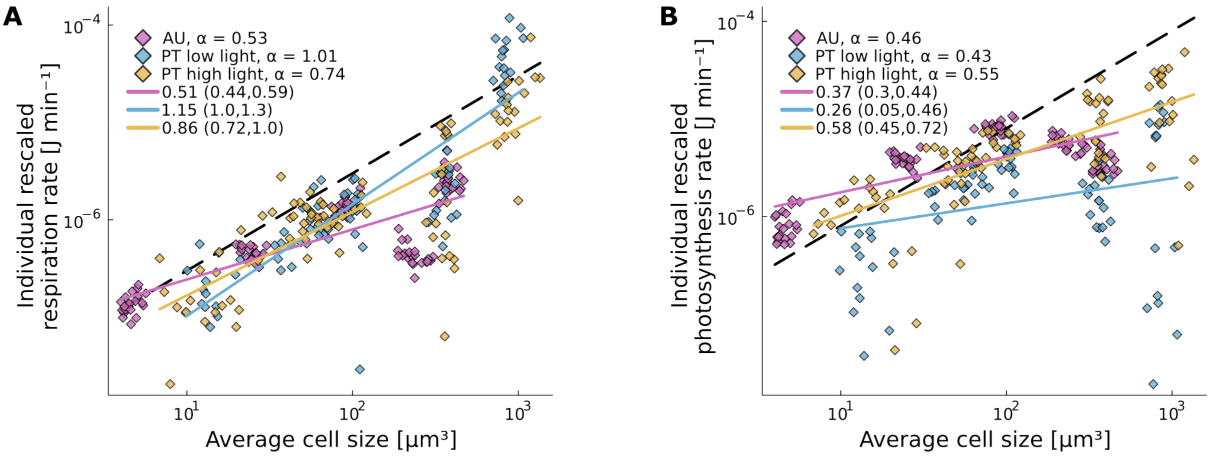


**Fig. S6. Scaling of *per capita* metabolism at fixed population density.**

*Per capita* respiration (**A**) and photosynthesis (**B**) scale allometrically with cell size at fixed population densities following scalings ($\alpha$) that are compatible with those observed at the population level according to equation 2 in the main text (and reported in the legend here).

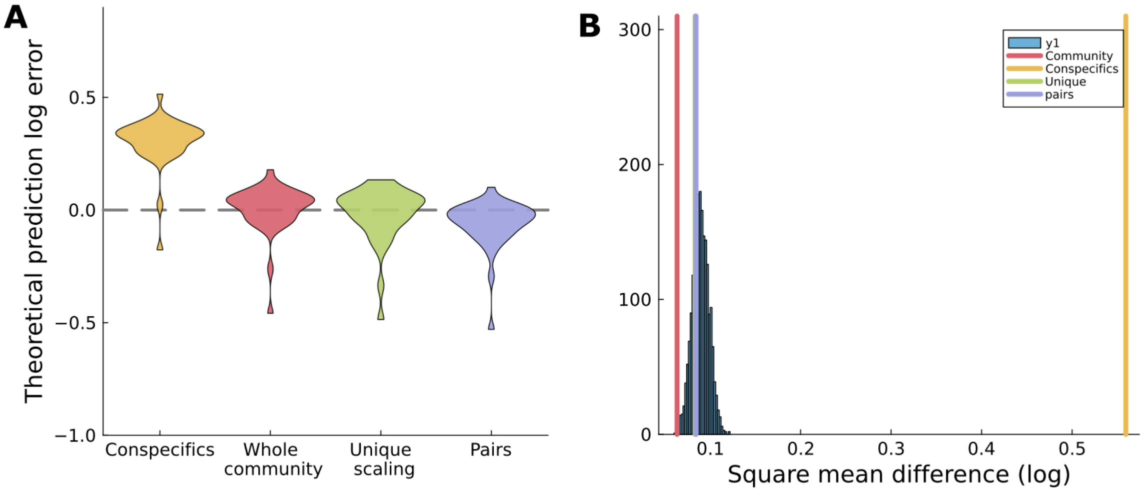


**Fig. S7. Predictions of community respiration based on the effects of each competitor on one another determined using pairs of species.**

We explored an intermediate situation in which interspecifics might affect metabolism (here shown for respiration) in a weaker way than conspecifics ("pairs", represented in purple). Despite containing more information (we account for changes in a species metabolism in response to each competitor), this approach does not perform significantly better than the approach based on responses to all competitors ("whole community", magenta) or the "unique" decline in metabolism with biomass across all species (green). (**A**) Shows the error of the prediction on log scale. (**B**) Shows that all approaches, except the one based on conspecific biomass only, fall within the distribution of predictions obtained from randomizing the association between species-specific declines in metabolism and species biomass in the community. This indicates that species identity does not have a strong effect on the metabolic decline we observe in communities with increasing biomass. Note that estimates based on pairs can only be calculated for one dataset (AU data) because they require data on each pairs of species.

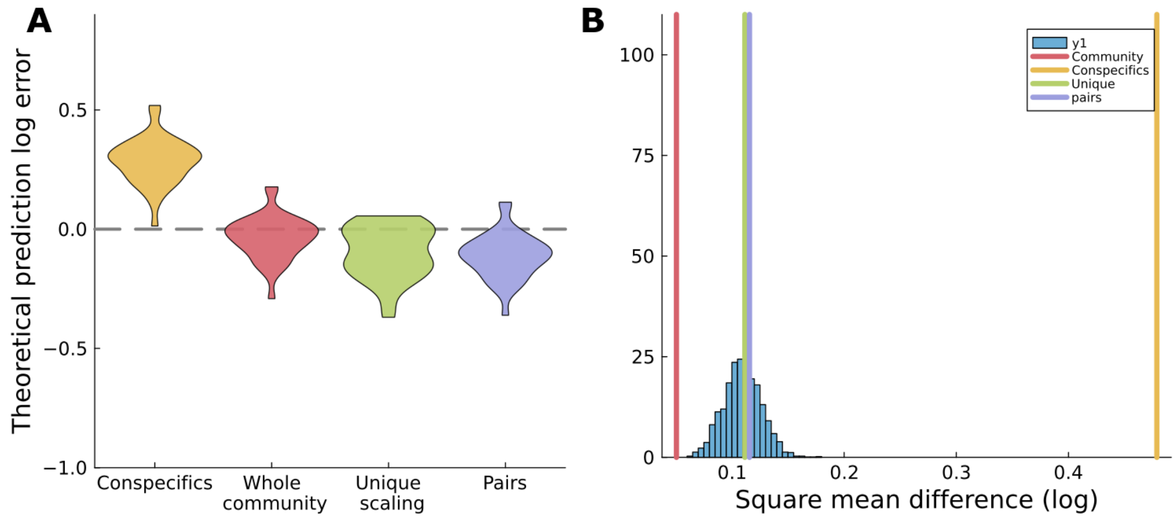


**Fig. S8**. **Predictions of community photosynthesis based on the effects of each competitor on one another determined using pairs of species.**

As Fig. S7 but predictions for photosynthesis based on pairwise responses.

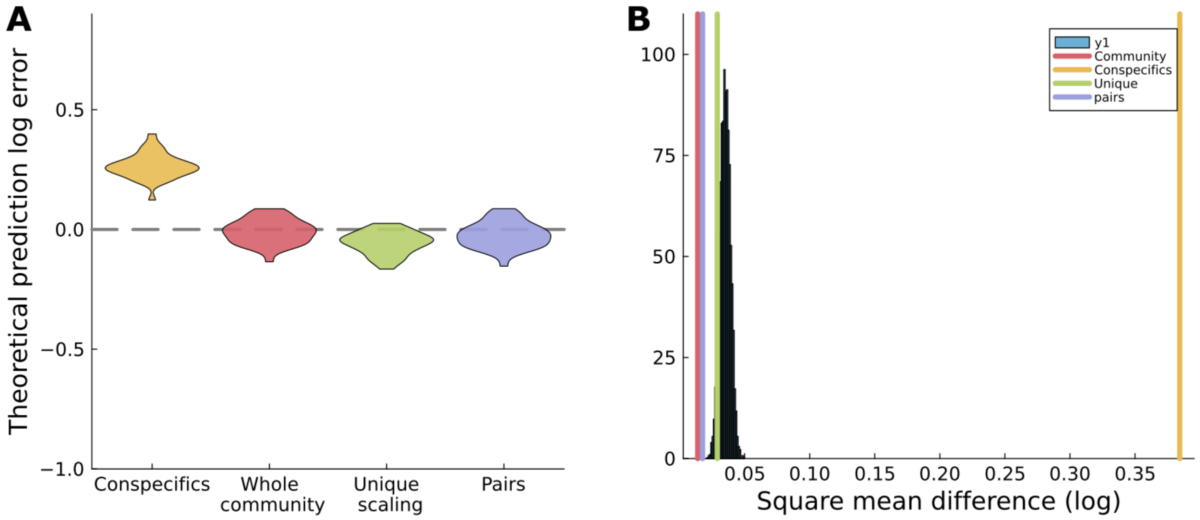


**Fig. S9. Predictions of community production based on the effects of each competitor on one another determined using pairs of species.**

As Fig. S7 but predictions for biomass production based on pairwise responses.

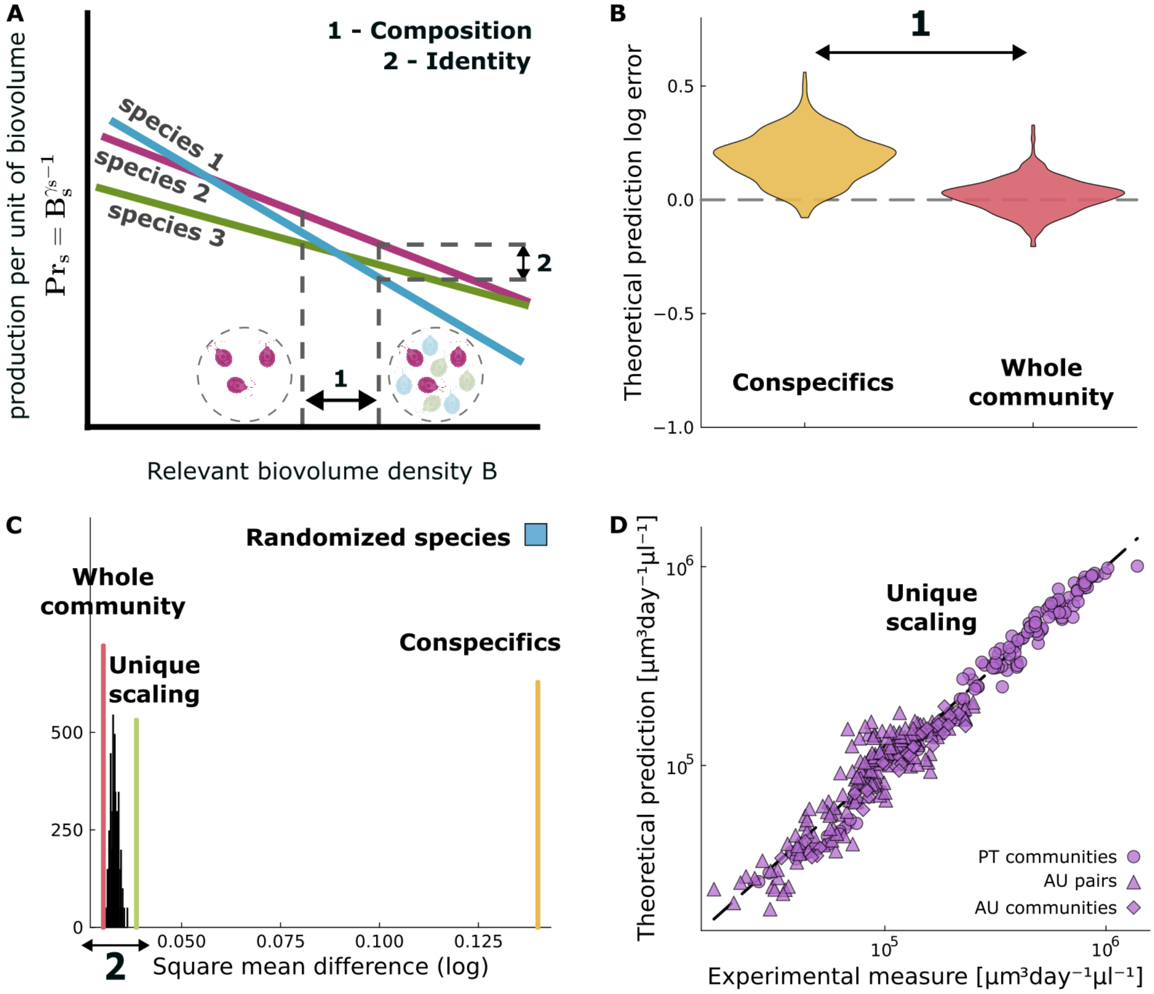


**Fig. S10. Predictions of community production based on monoculture data.**

(**A**) Biomass production per unit biovolume declines with increasing biovolume density; we use these relationships to test the relative importance of 1) *biovolume composition* (which competitors matter?) and 2) *species identity* on community production. (**B**) Only predictions based on total biovolume (accounting for declines in production in response to both intra- and inter-specific competitors) correctly predict community production, as we found for respiration (Fig. 3B) and photosynthesis (Fig. S11B). Estimates based on conspecifics overestimate community rates by underestimating density-dependence. (**C**) The "whole community" approach based on species-specific production rates performs better than a "unique" scaling across all species, but both fall very close to the distribution of randomized predicitons. Thus, while there are species-specific differences these seem to have minor effects on community productivity. (**D**) Indeed, the unique scaling between production and biomass density across species is sufficient to estimate production.

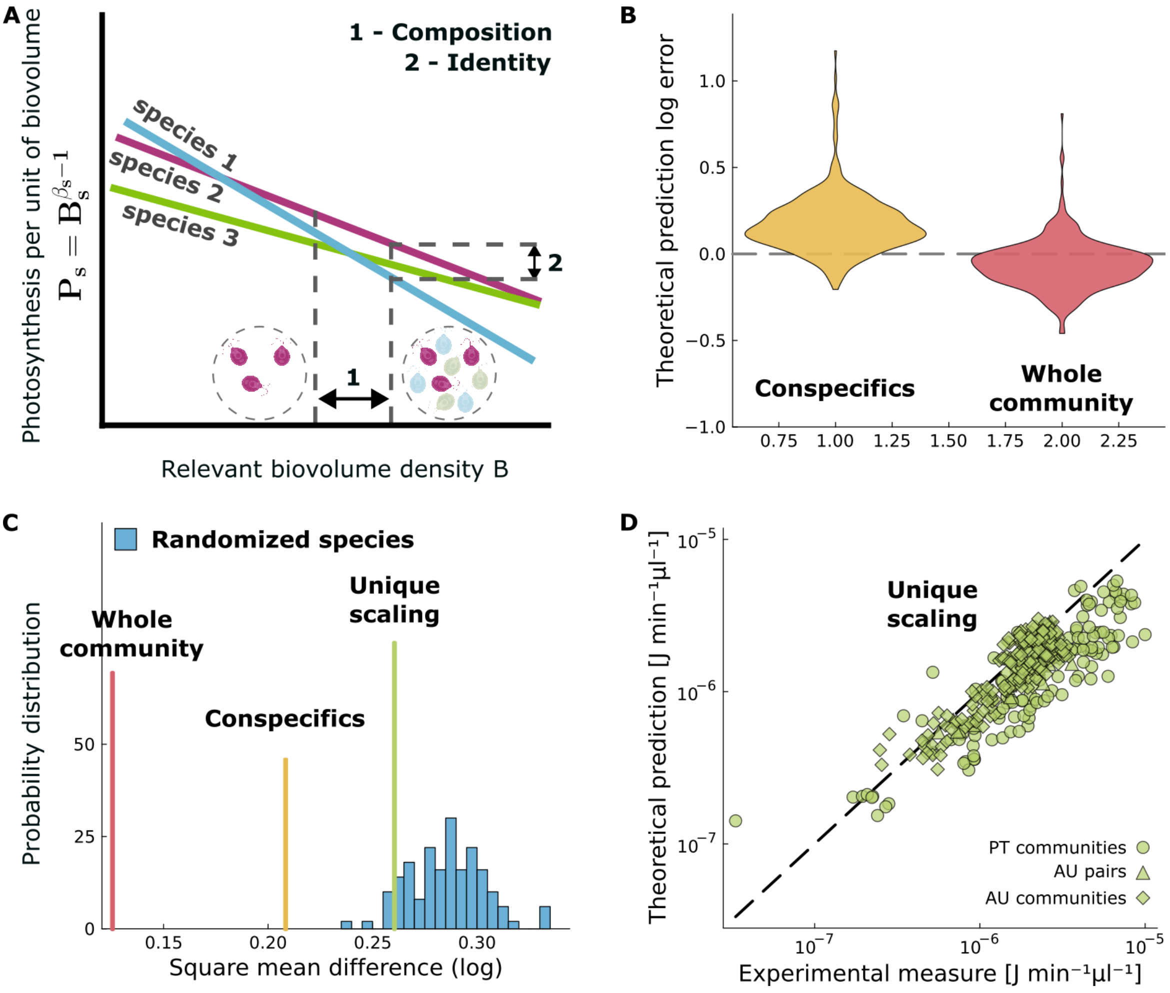


**Fig. S11. Predictions of community photosynthesis based on monoculture data.**

(**A**) We test the importance of (1) *biomass composition* and (2) species *metabolic identity* for predicting community photosynthesis. (**B**) As for respiration (Fig. 3B), predictions based on declines in photosynthesis per unit biovolume work well when based on total biomass, while those based on conspecifics biomass overestimate community rates. (**C**) But species identity has stronger effects on photosynthesis rates than on respiration (Fig. 3C), because we cannot predict community photosynthesis as well when randomizing or using a unique scaling across species (see also panel **D**).

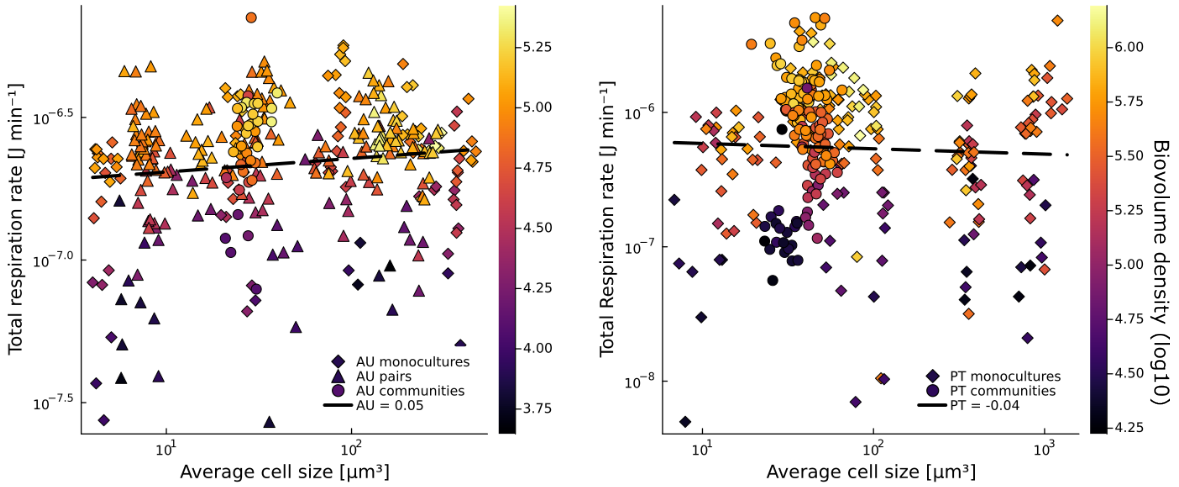


**Fig. S12. Total energy use is independent of body size but varies with total biomass.**

While the relationship between total energy use and body size is roughly around zero (reminiscent of energy equivalence), populations and communities composed of organisms of similar size can have different total rates of energy use (left panel: AU data; right panel: PT data). In our system, much of this variability is explained by biomass accumulation over time.

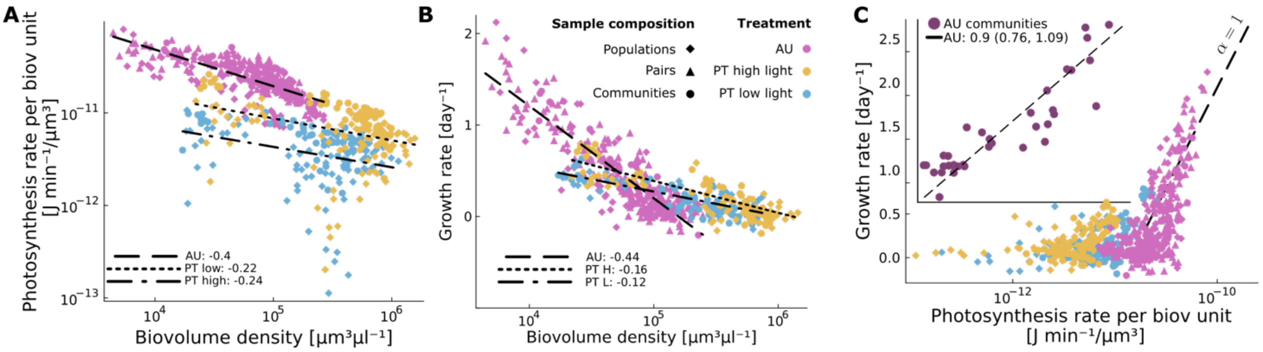


**Fig. S13. Relation between photosynthesis per unit biovolume and biomass production.**

(**A**) Photosynthesis per unit biovolume declines with increasing biovolume density at different rates depending on the environment. (**B**) This relationship is similar to the decline in biomass growth with total biomass but, conversely to respiration (Fig. 4), there is a weaker quantitative link between the two (**C**).
